# SAGERank: Inductive Learning of Protein-Protein Interaction from Antibody-Antigen Recognition using Graph Sample and Aggregate Networks Framework

**DOI:** 10.1101/2023.10.11.561985

**Authors:** Chuance Sun, Ganggang Bai, Honglin Xu, Yanjing Wang, Buyong Ma

## Abstract

Numerous experiments and computationally solved antibody-antigen interfaces offer the possibility of training deep-learning models to help predict their biological correlations. Predicting antibody-antigen docking and structure-based design represent significant long-term and therapeutically important challenges in computational biology. We present SAGERank, a general, configurable deep learning framework for antibody design using Graph Sample and Aggregate Networks, which mainly includes ranking docking decoys, detecting binding, and identifying biological interfaces. The model proved its reliability in three different tasks. For both problems ranking docking decoys and identifying biological interfaces, SAGERank is competitive with or outperforms, state-of-the-art methods. Besides, the SAGERank model still showed a high degree of confidence in determining whether the antibody-antigen could bind. All of these demonstrate the versatility of SAGERank for structural biology research. Most importantly, our study demonstrated the real potential of inductive deep learning to overcome small dataset problem in molecular science. The SAGERank models trained for antibody-antigen docking can be used to examine generally protein-protein interaction docking and differentiate crystal packing from biological interface.

## Introduction

The recognition of foreign antigens by antibodies is the crucial step in immune response, and deciphering antibody-protein antigen recognition is of fundamental and practical significance. The antibody-protein antigen interaction is a subset of general protein-protein interaction (PPI), and both categories share similar principles but antibody-antigen interaction involve distinctly different sequence and structural features from non-antibody PPI ^1,2^. Consequently, the general protein-protein docking programs, such as ZDOCK and HADDOCK, often need specialized treatment for their application in antibody-antigen docking. Specially benchmark or program for antibody-antigen docking^3, 4^ have been developed, including traditional docking method and recent AI approaches^5^. Overall, two major problems exist for computational study of antibody-antigen interaction. The first one is the relatively small dataset available and used for antibody-antigen training. For example, the expanded benchmark for antibody-antigen docking has only 67 antibody-antigen cases. The second question is a related but more general situation in modern machine learning training: how to obtain meaningful information from small data^6^.

The high expressive power of deep neural networks enables efficient training with a large amount of data^7^. Enlarging the antibody-antigen certainly is the right direction^8^, some approach also use a combination of structure modeling and computational docking to create training data set of antibody-antigen complexes. Still the problem stands: if we can use general protein-protein interaction data to study antibody-antigen recognition or vice versa? For the small data challenges in molecular science, exploration of the latest advances in deep learning algorithms and new methods are needed.

In the last few years, deep learning techniques have attracted much attention as a promising alternative to the physicochemical based approaches^9^. Compared to docking calculations, deep learning methods have improved the performance by learning the extracted features from protein-ligand complexes^10^. It can automatically extract task-related features directly from data without handcrafted features or rules. Among many deep learning methods, various graph neural networks (GNN) are especially suitable for the questions related to protein structure and protein-protein interactions, as illustrated in a recent Hierarchical Graph Neural Networks for Protein–Protein Interactions.^11^

Here, we applied a Graph Sample and Aggregate Network (called GraphSAGE) to the problem for ranking antibody-antigen docking models, detecting antibody antigen binding, and identifying biological and crystal interfaces. Low-dimensional vector embeddings of nodes in large graphs have proved extremely useful as feature inputs for a wide variety of prediction and graph analysis tasks^12,13,14,15,16^. However, most embedding frameworks are inherently transductive and can only generate embeddings for a single fixed graph. These transductive approaches do not efficiently generalize to unseen nodes (e.g., in evolving graphs), and these approaches cannot learn to generalize across different graphs^17^. In contrast, GraphSAGE is an inductive framework that leverages node attribute information to efficiently generate representations on previously unseen data. As far as we know, this is the first work to apply GraphSAGE to the ranking decoys, detecting binding and identifying interfaces about antibody-antigen. Our work demonstrated that the model trained for ranking antibody-antigen docking poses can directly applied to different tasks of distinguishing antibody antigen pairing, and even extending to non-antibody protein-protein interactions.

## Materials and Methods

### SAGERank architecture of antibody-antigen interfaces

SAGERank is built as a Python 3 package that allows end-to-end training on datasets of 3D Antibody-Antigen (Ab-Ag) complexes. Figure 1 shows the architecture of the network. The framework consists of two main parts, one focusing on data pre-processing and featurization and the other on the training, evaluation, and testing of the neural network.

**Fig. 1.**
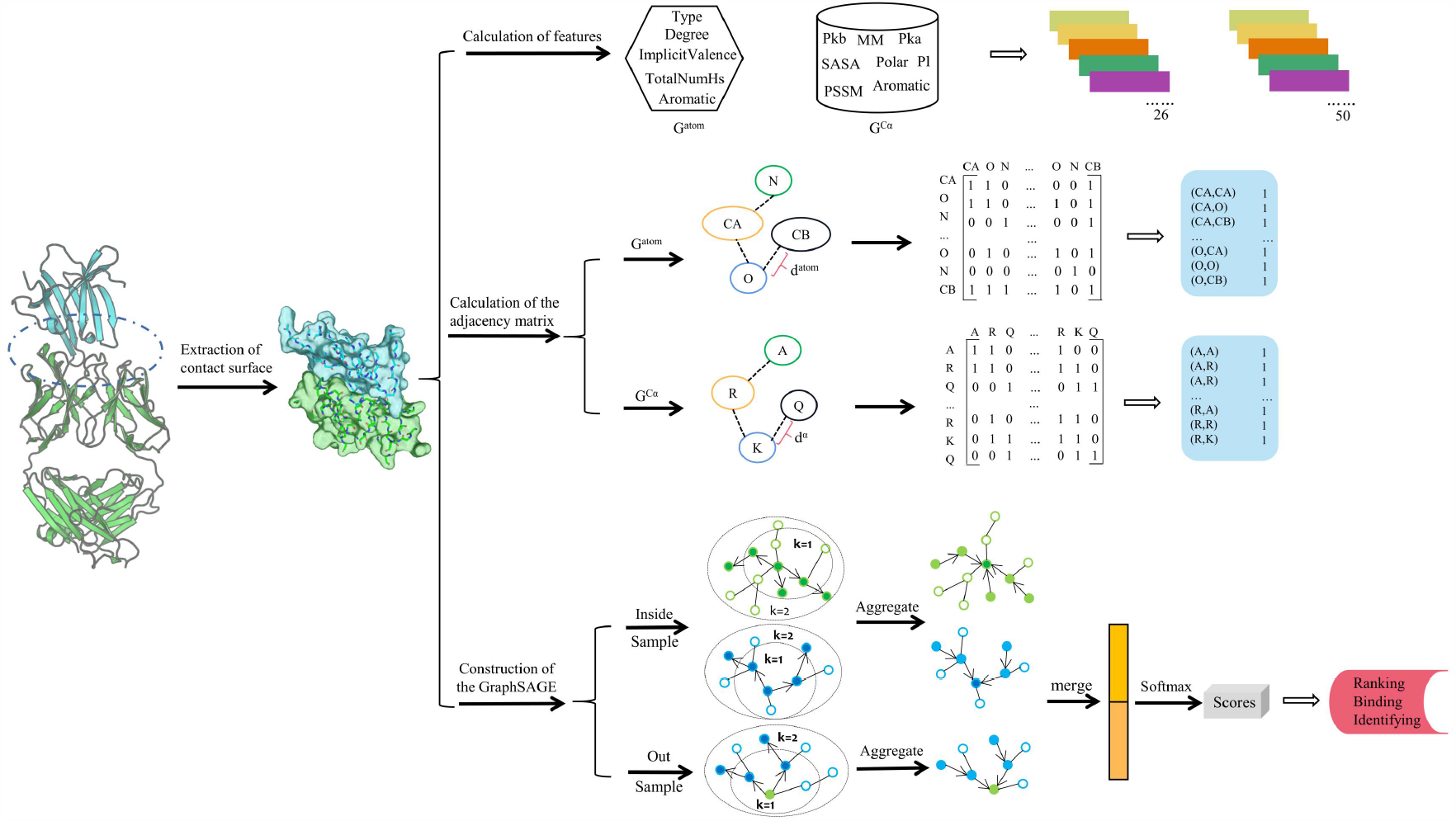
The framework of SAGERank. SAGERank extracts the interface region of antibody-antigen complex and further constructs graph based on atom nodes(26 features) and residue nodes(50 features). In order to save memory, we compress the sparse adjacency matrix by COO(Coordinate Format). The COO contains three arrays that store the row index, column index, and value of all non-zero elements. The graph represention of interface region is split into two sub-graphs that is internal graph (include receptor or ligand) and external graph (include receptor and ligand). The internal and external graph are sequentially passed to 4 consecutive SAGEConv layers and 1 global mean pooling layer. The two final graph representations are merged before applying softmax function for output.

We have considered two types of networks, one based on amino acid represented by Ca coordinates (G^Ca^) and another by specific atoms in each amino acid (G^atom^). Starting from the 3D structures of Ab-Ag complexes and the interface region is identified as a set of residues located within 10.0 Å of any residues of the other antibody or antigen (Fig. 1). Graph G is defined by V (node set), E (set of edges), and A (adjacency matrix). Therefore, the amino acids in the interface region and the atoms that make up the amino acids represent two different nodes. The residue-based and atomic node features are listed in Table 1 and Table 2, respectively. Overall, 50-bit features and 28-bit features are used to describe the residue nodes and atom nodes, respectively.

**Table 1.** Features computed in residue network G^Ca^.

| Name of the feature | Full name | Description | Dimension |
| --- | --- | --- | --- |
| Type | Residue type | One-hot encoded | 20 |
| Aliphatic | Residue aliphatic | One-hot encoded | 1 |
| Aromatic | Residue aromatic | One-hot encoded | 1 |
| Polar neutral | Residue polar neutral | One-hot encoded | 1 |
| Charged | Acidic/Basic charged | - | 2 |
| Weight | Residue weight | - | 1 |
| Pka | The negative of the logarithm of the dissociation constant for the $-COOH$ group | - | 1 |
| Pkb | The negative of the logarithm of the dissociation constant for the $-NH_3$ group | - | 1 |
| Isoelectric point | Residue isoelectric point | - | 1 |
| Hydrophobicity | Hydrophobicity of residue (pH=2) | - | 1 |
| Hydrophobicity | Hydrophobicity of residue (pH=7) | - | 1 |
| PSSM | Position-specific scoring matrix | - | 20 |
| SASA | solvent-accessibility surface area | - | 1 |

**Table 2.** Features computed in atomic network G^atom^.

| Name of the feature | Full name | Description | Dimension |
| --- | --- | --- | --- |
| Type | Atom type | One-hot encoded | 10 |
| Degree | Number of atomic connections | One-hot encoded | 6 |
| TotalNumHs | Number of hydrogen atoms | One-hot encoded | 5 |
| ImplicitValence | Atomic implicit valence | One-hot encoded | 6 |
| Aromatic | Atomic aromatic | - | 1 |

Adjacency matrix is constructed by the following procedure. For amino acid network G^Ca^ with N nodes, the adjacency matrix A^Ca^ has a dimension of N*N (eqn 1). Within the ligand (antigen) and receptor (antibody) where A^Ca^ _ij_ =0 if the Euclidean distance d^Ca^ _ij_ between i-th node and the j-th node is greater than 4.5Å, and A^Ca^_ij_ =1 otherwise. In addition, we also take into account the specific properties of the antibody CDR region in the receptor where A^Ca^_ij_ =1 if i-th node and the j-th node belong to the CDR loops, and A^Ca^ _ij_ =0 otherwise. Between receptor and ligand where A^Ca^_ij_ =1 if the distance d^Ca^_ij_ between i-th node and the j-th node is greater than 10Å, and A^Ca^ _ij_ =0 otherwise. For atomic network G^atom^ with M nodes, the adjacency matrix A^atom^ has a dimension of M*M (eqn 2). Within the receptor and ligand where A^atom^_ij_ =1 if atom i and atom j are connected by a covalent bond, and A^atom^_ij_=0 otherwise. Between receptor and ligand where A^atom^_ij_ =1 if the distance d^atom^_ij_ between i-th node and the j-th node is greater than 8Å, and A^atom^_ij_=0 otherwise.

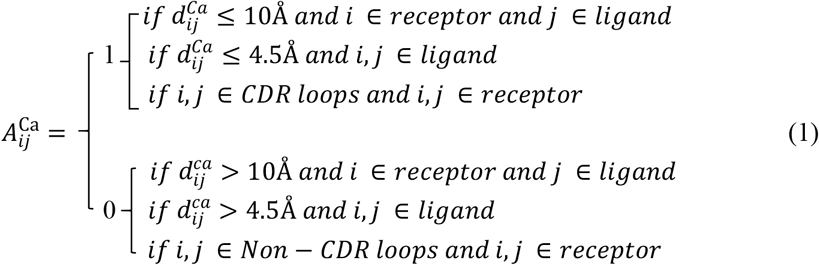

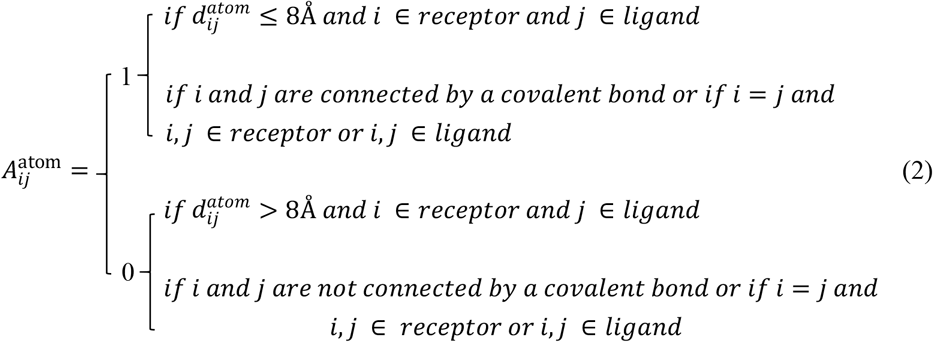

Node aggregation and update follow the standard GraphSAGE algorithm. The core steps of GraphSAGE are neighbor sampling and feature aggregation. The forward propagation algorithm for GraphSAGE is as follows:

#### GraphSAGE emdedding generation algorithm

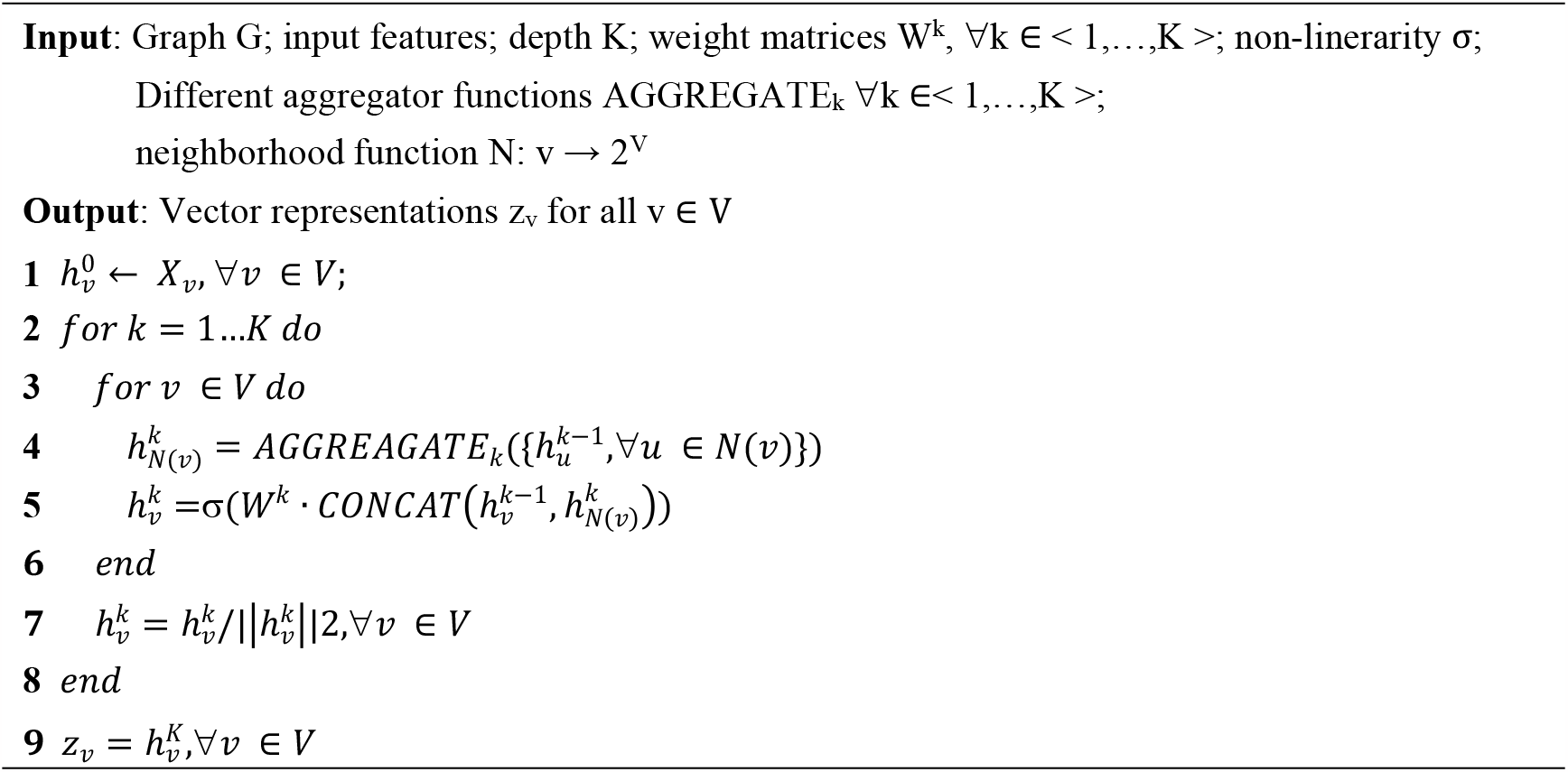

In the algorithm, the first for loop is used to traverse the number of layers, and the second for loop is used to traverse all nodes in the Graph. Sampling is performed on the neighbors of each node ν to obtain N ν. Next, aggregate the embedding of neighbor nodes through AGGEGATEk to obtain 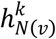 Then 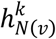 is spliced with the current embedding 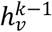of the target node, and assigned to 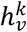 after nonlinear transformation, thereby completing an update of the target node ν. When the for loo (*k* = 1…*K*) traversal of the outer layer ends, the node ν will complete the information aggregation of k-order neighbors.

### Training hyperparameters and quality metrics

The training hyperparameter settings are shown in Table 3. The entire method is implemented using the Pytorch-geometric deep learning library.

**Table 3.** The hyperparameter settings using human experience.

| Hyperparameter | Setting |
| --- | --- |
| Epoch | 50 |
| Batchsize | 512 |
| Learning rate | 0.0003 |
| Optimizer | Adam |
| Loss function | CrossEntropyLoss |
| The layers of GraphSAGE | 4 |

Hit-rate and Success-rate are used to evaluate the performance of different scoring functions for ranking docked decoys. The Hit-rate is defined as the percentage of near-native (models with iRMSD ≤4 Å) models in the top-ranked models for a specific complex:

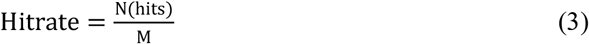

where N(hits) is the number of near-native models among top models and M the total number of near-native models for this case. The Hit-rate was calculated for each individual case in our test set and the higher the value, the higher the accuracy of ranking decoys. The Success-rate is defined as the percentage of complexes for which at least one near-native model is found in the top K selected models. It is therefore defined as:

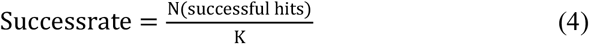

where N(successful hits) is the number of cases with at least one near-native model among top models, and K is the total number of cases. The commonly used statistical indicators for measuring binary classification problems are Accuracy (eqn 5), Precision (eqn 6), Recall (eqn 7) and F1 (eqn 8). Among them, TN is true negative, TP is true positive, FN is false negative and FP is false positive. When the number of positive and negative samples in the dataset is relatively balanced, accuracy can be used as an evaluation indicator, but when the number of positive and negative samples is unbalanced, F1 values and ROC are more trustworthy. The ROC curve is defined as the fraction of the true positive rate as a function of the fraction false positive rate while navigating through the ranking provided by the scoring function. The AUC is the integral of the ROC curve and is equal to 1 for an ideal classifier and 0.5 for a random classifier.

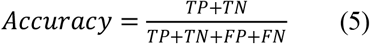

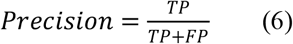

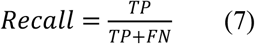

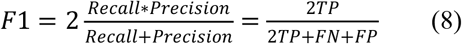

## Results

### 1. Training and performance of SAGERank docking model to rank antibody antigen docking poses

We have collected a dataset of 287 different Ab-Ag complexes with sequence identity < 95%. We used megadock^18^ to generate a set of docking poses of various RMSDs for all Ab-Ag complexes. In the process of docking, we adopt a semi-flexible docking method, that is, the receptor conformation does not change, and the ligand rotates in the CDR region of the antibody. 20,000 different conformations were generated for each Ab-Ag complex. Therefore, based on interface iRMSD, we divided the decoys of docking into two categories: the ones with iRMSD greater than 4 Å are negative models, and those with iRMSD less than 4 Å are near-native poses. In the end, 287 different Ab-Ag complexes generated a total of 455,420 docking decoys with 1:3 ratio of positive and negative samples. The conformations were divided into training (80%), validation (10%), and test (10%) sets. In addition, in order to compare with other docking methods, we constructed 10 groups of Ab-Ag complexes and 8 protein-protein complexes with 23,508 and 14,278 docking decoys, respectively.

As can be seen in Supplementary Fig.1, the model with atoms as nodes achieves much better ranking accuracy than amino acids as nodes. There are two factors contributing to the different performances. Firstly, the graph core and the interaction between interfaces can be more accurately represented and captured using a larger number of nodes of atomic network G^atom^ than residue based network G^Cα^. Secondly, the atomic network G^atom^ catches essential physicochemical features underlying antibody-antigen recognition.

We then compared SAGERank with four leading scoring functions in ranking protein-protein docking poses: Zrank^19^, Pisa^20^, FoldX^21^, and Rosetta^22^ (Fig.2B, Supplementary Fig.2 and Supplementary Table.1). Clearly, SAGERank performs better than the other methods in the Ab-Ag docking decoys set. An analysis of success rate of SAGERank and Pisa in the 7MLH case (Supplementary Fig.3) confirms the good performance of SAGERank. In this case, if we select the consensus hits from top200 poses of SAGERank and Pisa, only a few false positive hits appear.

**Fig. 2.**
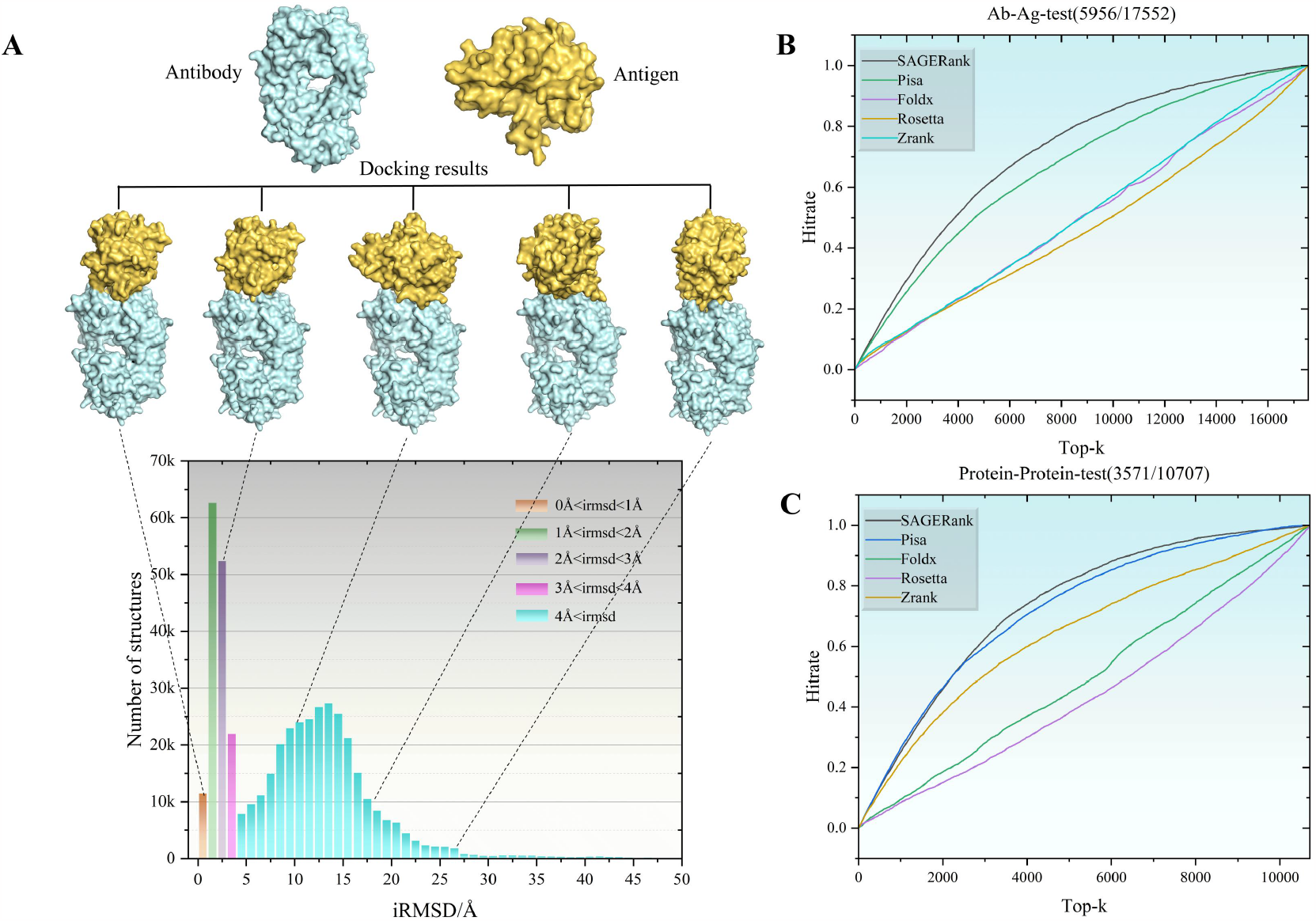
SAGERank applied to the ranking docking decoys problem. A: iRMSD distribution of all docking decoys in the training set. B: Performance of five different evaluation methods on Ab-Ag docking decoys dataset. C: Performance of five different evaluation methods on protein-protein docking decoys dataset.

Unlike common practice to use protein-protein interaction model for antibody-antigen docking, we test an opposite application of the SAGERank: use the model trained with small antibody-antigen dataset for general protein-protein dock rank. We constructed the protein-protein docking decoys set using 8 protein-protein complexes and 14,278 docking decoys. Surprisingly, we found that SAGERank is still competitive with these scoring functions specifically optimized for protein-protein docking, slightly outperforming Pisa (Fig2.C, Supplementary Fig.3 and Supplementary Table.2). Our results suggest the ability of SAGERank to correctly rank the docking decoys, whether Ab-Ag complex or protein-protein complex.

### 2. SAGERank model2: Predicting if antibody can bind to an antigen in right epitopes

A dataset with shuffled antibody-antigen pairing was generated using the megadock. Firstly, the sequence alignment of 287 different antigen-antibody complexes was performed to remove the entries with similarity between antigens > 70%, and the sequence similarity between antibodies > 95%. After that, we obtained 230 Ab-Ag complexes of which 200 complexes were used as the training set and the remaining 30 complexes were used as the test set. Next, the positive samples consist of the decoys of the iRMSD in the 3.5 Å range generated by docking the antibody with its own antigen. The negative sample dataset consists of two parts: one is decoys with iRMSD greater than 8Å generated by docking between the antibody and its own antigen, and the other is the structure generated by docking between antibodies and the remaining 199 antigens. In the end, the dataset produced a total of 369,932 complex structures, with a ratio of positive and negative samples of about 1:5. Among them, 80% is used as the training set, 10% as the verification set, and 10% as the test set.

Because the proportion of positive and negative samples in the dataset is not balanced, in this situation, it is more reliable to choose Auc for model evaluation in this case, with Auc reaching 0.82 in the test set (Supplementary Figure 6), slightly smaller than the AUC obtained in docking pose ranking model. Next, we selected the optimal threshold (0.3) based on the changes in F1-score under different thresholds (Supplementary Figure 7). F1-score is an indicator used in statistics to measure the accuracy of binary classification models. It takes into account both the accuracy and recall of the classification model. Fig.3.B is the distribution of SAGERank scores for positive and negative samples in test sets. It is obvious that the scoring values of most negative samples are concentrated below 0.3. The scores of positive samples are mostly above 0.3 and more evenly distributed. Therefore, selecting 0.3 as the threshold can maximize the performance of the model.

**Fig. 3.**
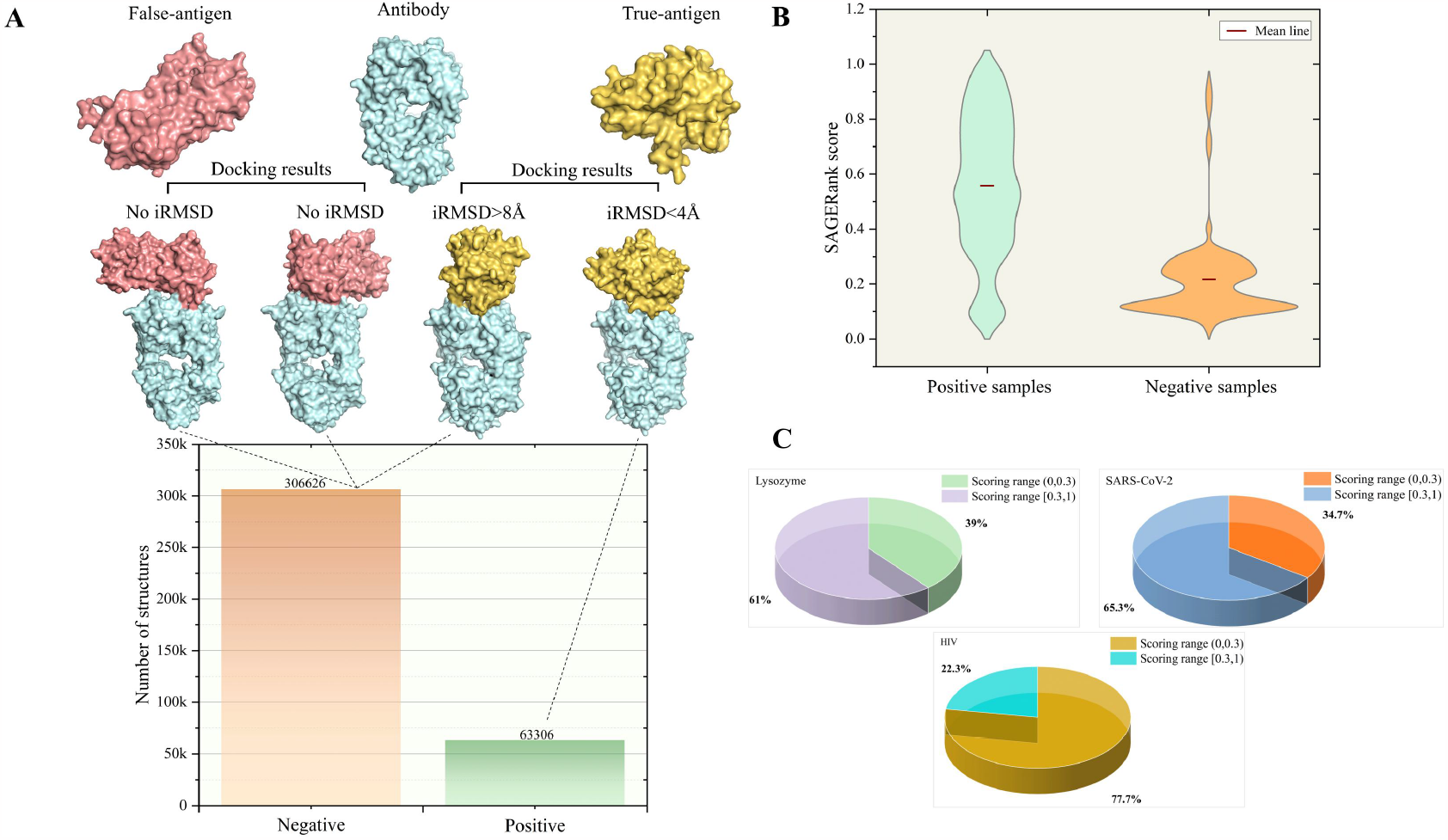
SAGERank applied detecting antibody antigen binding problem. A: The number of complexes on the training set where antibodies bind to antigens (positive samples) and complexes where antibodies cannot bind to antigens (negative samples). B: SAGERank’s scoring distribution for positive and negative samples on the test set. C: Recognition success rate of SAGERank for antibody antigen complexes of SARS-CoV-2, HIV and Lysozyme.

**Fig. 4.**
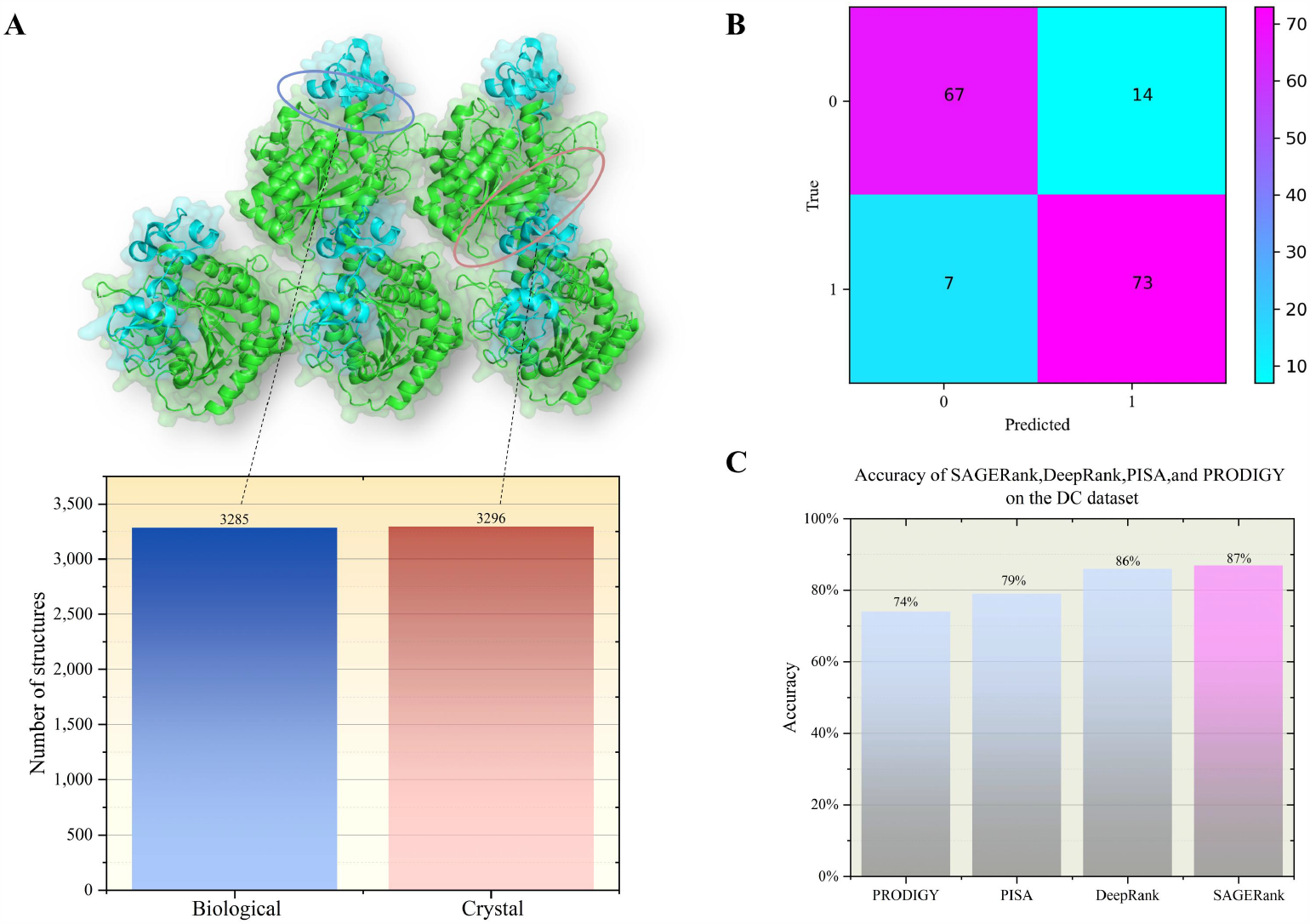
SAGERank applied identifying biological and crystal interfaces problem. A: The number distribution of biological and crystal interfaces on the training set. B: The confusion matrix of SAGERank on the DC test dataset. C: Comparison of accuracy of four different methods on DC data sets.

We further tested the model to examine antibody-antigen complexes for three targets with large number of structures available, including 222 structures for the SARS-CoV-2 target, 41 structures for the Lysozyme target, and 367 structures for the HIV target. In other words, 630 positive samples were chosen to test the distinguishing ability of SAGERank model2. From the Fig.3.C we can observe that the recognition success rates of the model for three targets HIV, SARS-CoV-2 and Lysozyme are 77.7%, 65.3%, and 61.0%, respectively. Based on the above analysis and validation, we believe that the SAGERank model2 has achieved a considerable success rate in determining whether antibodies and antigens can bind. It is interesting to comparing the performance of the SAGERank docking model trained with the antibody-antigen docking poses. The success rates of the docking model for three targets SARS-CoV-2, HIV, and Lysozyme are 50%, 48.6%, and 70.0%, respectively. It is worth noting that the recognition accuracy of the docking model for antigens and antibodies in lysozyme group is higher than that of the specifically trained SAGERank model 2.

### 3. SAGERank model3: Identifying biological and crystal interfaces

Over 80% of the protein structures in the PDB library are obtained through X-ray crystallography methods^23^. The molecular packing in crystals contain multiple interfaces, some of which are truly biological recognitions, while others are just packing contact in the crystal, known as the “crystal interface”. The commonly used DC dataset contains 80 biological and 81 crystal interfaces. At the moment, DeepRank^23^, PISA^24^, and PRODIGY-crystal^25, 26^ show the highest prediction performances in distinguishing crystal interfaces from biological ones.

In addition to the crystal and biological interface datasets (https://github.com/eppic-team/datasets/blob/docking/data), we added more entries from our Ab-Ag datasets. Using the eppic WebServer (https://www.eppic-web.org/), we extracted protein-protein complex structures with 4001 biological interfaces and 4153 crystal interfaces. In the training, 80% is used as the training set, 10% as the verification set, and 10% as the test set.

We first test if the SAGERank docking model trained for antibody-antigen docking pose ranking can be used to distinguish the crystal and biological interfaces in the DC dataset. We found that SAGERank docking model has a 79.5% success rate in telling biological interface from crystal interface.

We then retrain the model specifically using the crystal/biological interface dataset. During the training period, the model demonstrated excellent performance in the test set, with accuracy and AUC reaching 0.9 and 0.95, respectively (Supplementary Fig.9 and Supplementary Fig. 10), much higher than that for models 1 and 2. Besides, to compare with other published methods, we examine the performances using the DC dataset. SAGERank correctly classified 73 out of 80 biological interfaces and 67 out of 81 crystal interfaces (Fig.3.B). SAGERank achieved an accuracy of 87%, outperforming PRODIGY, PISA, and DeepRank, which reported 74%, 79%, and 86%, respectively. Note that the training set of the SAGERank also includes a portion of the interface structure of antibodies. In contrast, the training set in Deeprank only includes the interface structure of proteins. This demonstrates that SAGERank can accurately distinguish between most biological and crystal interfaces for proteins and antibodies.

## Discussion and Conclusion

High-resolution structures of Ab-Ag complexes are necessary for understanding mechanisms of Ab-Ag interactions, analyzing mutations, and modulating binding affinity^27^. The large gap between the number of experimentally determined complex structures and the available sequences of pairs of Ab-Ag complexes underscores the challenges, time required, and cost of experimental approaches^28^. The paucity in complex structures can be alleviated by computational docking, which potentially provides a fast and efficient alternative route. protein docking methods that use unbound or modeled component structures as input to perform rigid-body global searches in six dimensions^29,30,31,32^, and template-based modeling methods that generate models of complexes based on known structure^33, 34^. Challenges for docking algorithms include side chain and backbone conformational changes between unbound and bound structures, large search spaces, and inability to capture key energetic features in grid-based and other rapidly computable functions, leading to false positive models among top-ranked models or lack of any near-native models within large sets of predicted models^35^. Although substantial improvements have been made in protein docking, selecting near-native models out of a large number of produced models is still challenging^36^.

Two major factors are responsible for the above difficulties. Firstly, while the principles of protein-protein interactions have been actively investigated during last two decades^2^, we also increasingly realize the complexity of protein-protein interaction. Many PTMs for example, phosphorylation and glycans exist and modulating protein-protein interfaces and interaction. Protein structure and conformation are dynamic^37, 38, 39, 40^, which added into hidden dynamical element in protein-protein interaction. The dynamics is especially important for antibody-antigen recognition due to the highly flexible CDR regions in antibody.

Still recently advances in protein structure prediction, represented by the successes of Alphafold2 and Alphafold-multimer greatly boosted the accuracy of predictions of protein-protein interaction and antibody-antigen recognition. Still, the traditional transductive learning limited the application of proteins with unusual sequence features and those with only small data available, which is the second major difficulty need to be solved for the development of biological drugs.

Here we proposed SAGERank framework to address the above critical challenges. Overall, our model has demonstrated excellent performance in all three applications. First of all, in the application1 of ranking docking decoys, SAGERank outperformed major existing scoring functions compared. A natural graph network with atoms as nodes can be formed in the interface between antibody-antigen complexes and protein-protein complexes to accurately capture the fundamental physicochemical features between amino acid interactions. SAGERank can also accurately distinguish between the dataset of antibodies that bind to real antigens and false antigens (model2). Among them, the recognition success rate of antibody antigen complex of HIV target is close to 80%. Last but not least, when applied SAGERank to the classification of biological versus crystal interfaces (model3), it shows outstanding classification performance, better than other competing methods.

Most importantly, our study demonstrated the real potential of inductive deep learning, coupled with atomic interaction features, to overcome small dataset problem in molecular science. The SAGERank models trained for antibody-antigen docking can be used to examine generally protein-protein interaction docking and differentiate crystal packing from biological interface. In summary, we have designed a reliable and efficient deep learning framework for accelerating research based on antibody-antigen 3D structures, with the potential to expand to general protein-protein interaction. In the future, we will examine and models trained with larger dataset to fully explore the SAGERank’s potential.

## Supporting information

SAGERank-Supplementary Information

## Acknowledgments

The computational simulations are supported by the Center for High Performance Computing at Shanghai Jiao Tong University.

