## Supplementary material for "SAGERank: Inductive Learning of Protein-Protein Interaction from Antibody-Antigen Recognition using Graph Sample and Aggregate Networks Framework": SAGERank-Supplementary Information

### Supplementary Note 1: Ranking antibody antigen docking models

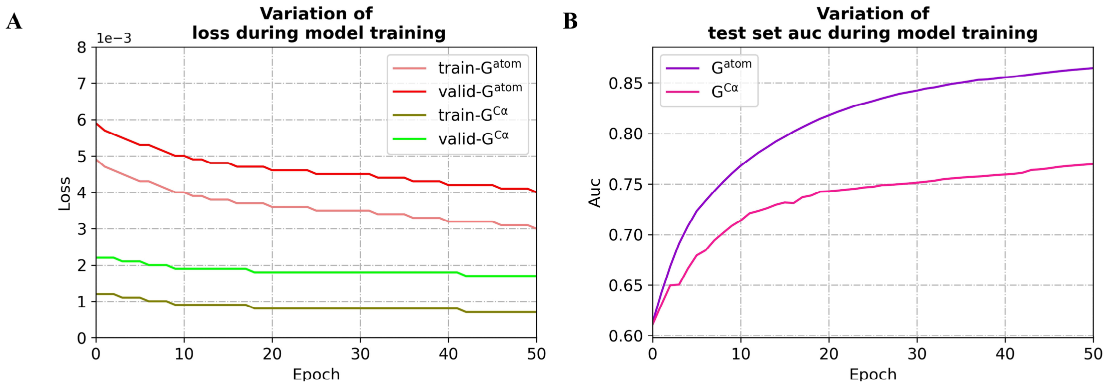


Supplementary Figure 1. Changes in loss (A) and AUC (B) during model training.

Supplementary Figure 1 shows the changes in loss and AUC during model training, from the Fig1.B we can observe that $G^{atom}$ with atoms as nodes have higher ROC values than $G^{C\alpha}$ with residues as nodes, this means it has more precise ranking accuracy. In this situation, in the following applications 2 and 3, we adopted similar graph structure construction methods.

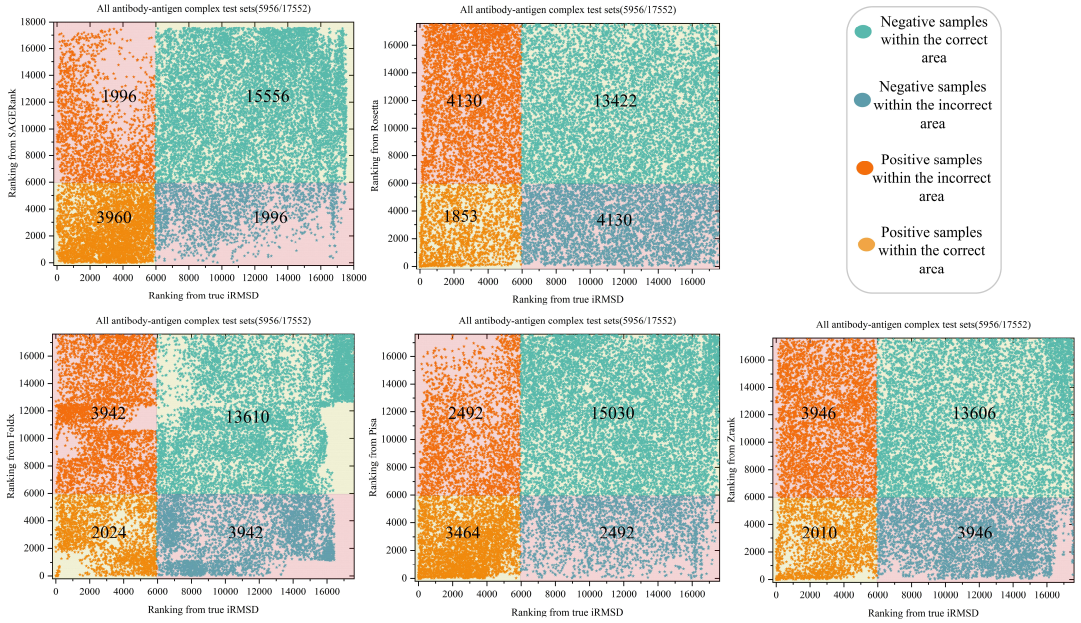


Supplementary Figure 2. Comparison of five different docking scoring methods for Ab-Ag docking decoys

In the Supplementary Figure 2, the abscissa is the ranking of all Ab-Ag docking decoys by iRMSD, and the ordinate is the ranking by SAGERank, Rosetta, Foldx, Pisa and Zrank, respectively. In this situation, we can notice that the SAGERank has the highest number of positive samples and negative samples within the correct area. At the same time, Pisa performed second, while Rosetta performed worst.

Supplementary Table 1. Performances on 10 cases of the Ab-Ag docking decoys set for five methods

| | Models | SAGERank | | | Pisa | | | Foldx | | | Rosetta | | | Zrank | | |
|---|---|---|---|---|---|---|---|---|---|---|---|---|---|---|---|---|
| | Native/ Total | T50 | T100 | T200 | T50 | T100 | T200 | T50 | T100 | T200 | T50 | T100 | T200 | T50 | T100 | T200 |
| 1W72 | 568/1704 | 34 | 77 | 164 | 33 | 72 | 138 | 33 | 58 | 102 | 26 | 41 | 70 | 6 | 28 | 79 |
| 3SM5 | 1630/4886 | 50 | 99 | 194 | 48 | 94 | 193 | 29 | 51 | 105 | 26 | 43 | 73 | 11 | 28 | 36 |
| 5D96 | 185/515 | 46 | 91 | 151 | 45 | 81 | 133 | 38 | 57 | 92 | 35 | 52 | 88 | 47 | 94 | 152 |
| 6WHK | 583/1709 | 40 | 87 | 171 | 43 | 83 | 162 | 22 | 48 | 96 | 27 | 44 | 82 | 19 | 42 | 98 |
| 6XXV | 1160/3440 | 5 | 9 | 40 | 7 | 12 | 44 | 22 | 48 | 92 | 48 | 78 | 127 | 1 | 4 | 16 |
| 7JWG | 122/326 | 24 | 57 | 104 | 45 | 65 | 86 | 7 | 25 | 75 | 27 | 53 | 93 | 30 | 51 | 89 |
| 7KQ7 | 715/2113 | 37 | 79 | 157 | 42 | 78 | 145 | 24 | 41 | 77 | 15 | 21 | 35 | 47 | 91 | 170 |
| 7MLH | 319/917 | 40 | 78 | 160 | 23 | 43 | 69 | 26 | 44 | 80 | 26 | 40 | 73 | 24 | 34 | 69 |
| 7VAE | 587/1721 | 43 | 88 | 178 | 24 | 55 | 113 | 1 | 8 | 24 | 24 | 42 | 66 | 0 | 0 | 0 |
| 7X8P | 87/221 | 41 | 69 | 87 | 48 | 81 | 87 | 25 | 43 | 74 | 21 | 38 | 71 | 23 | 42 | 76 |
| SR | - | 72% | 73% | 70% | 72% | 66% | 59% | 45% | 42% | 41% | 55% | 45% | 39% | 42% | 41% | 40% |

The number of near-native models among Top50, Top100 and Top200 are reported (Supplementary Table 1), showing the good ranking performance of SAGERank in this task. The bottom row (SR) indicates the overall success rate. From the **table** we can see that the SAGERank and Pisa achieved same SR (72%) in the Top50, and in the Top100 and Top200 SAGERank leads all other methods.

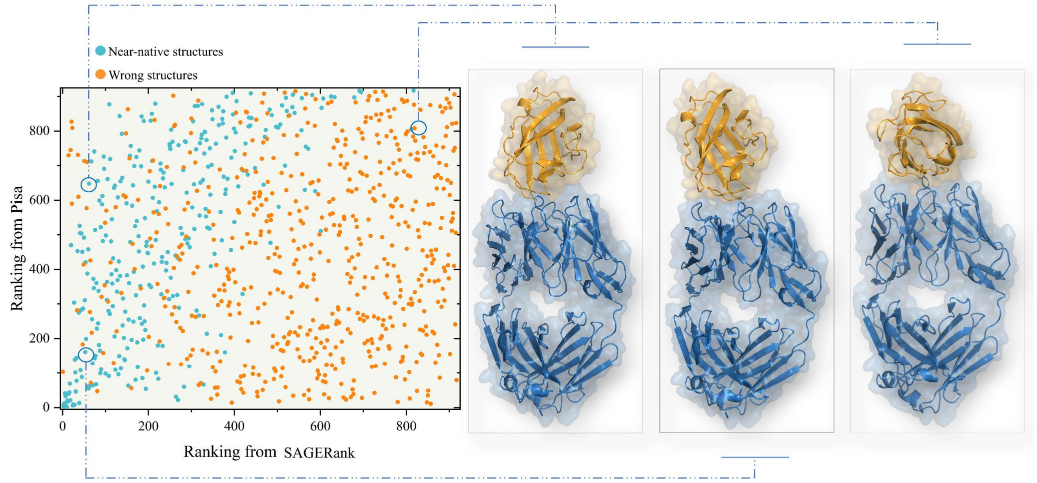


Supplementary Figure 3. Comparison of the rankings given by SAGERank and Pisa for the conformations of the 7MLH case

In the Supplementary Figure 3, the abscissa is the ranking of 7MLH test sets by SAGERank, and the ordinate is the ranking of test sets by Pisa. The blue/orange dots mark near-native/wrong models. This scatter plot illustrates near-native structures that are correctly or incorrectly ranked by either or both of the SAGERank and Pisa.

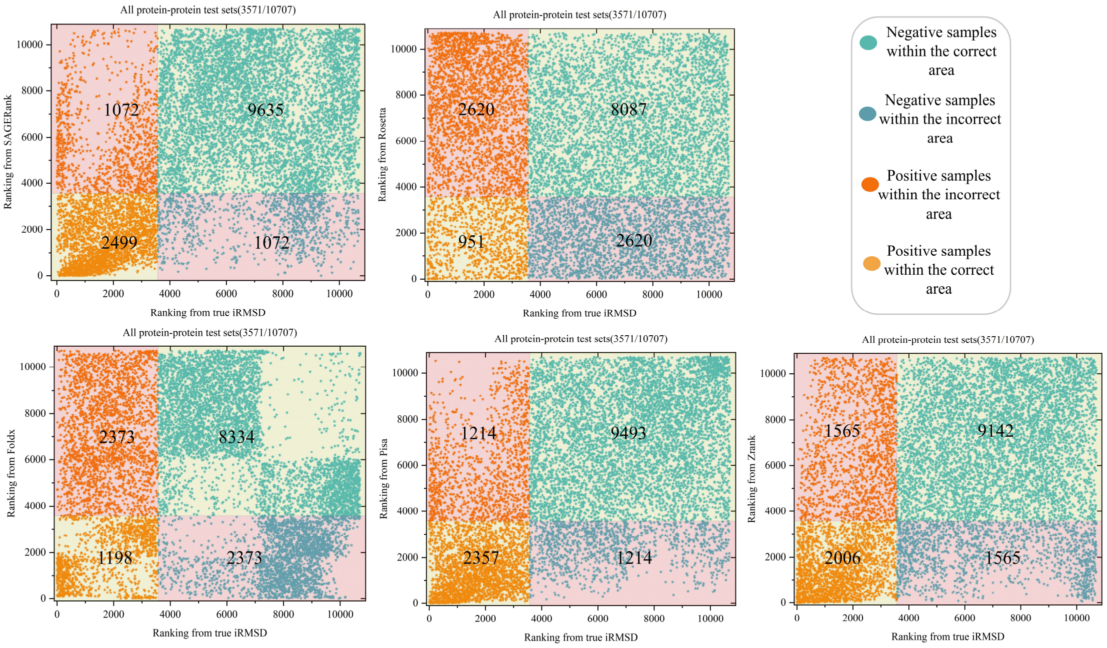


Supplementary Figure 4. Comparison of five different docking scoring methods for protein-protein docking decoys

The training set does not include the structure of protein-protein complexes, therefore, we would like to know if our model SAGERank can still maintain a high ranking accuracy of protein-protein docking decoys. According the Supplementary Figure 4, it is obvious that the accuracy of all protein-protein docking decoys for SAGERank still outperforms other methods.

Supplementary Table 2. Performances on 8 cases of the protein-protein docking decoys set for five methods

| | Models | SAGERank | | | Pisa | | | Foldx | | | Rosetta | | | Zrank | | |
|---|---|---|---|---|---|---|---|---|---|---|---|---|---|---|---|---|
| | Native/ Total | T50 | T100 | T200 | T50 | T100 | T200 | T50 | T100 | T200 | T50 | T100 | T200 | T50 | T100 | T200 |
| 1CGI | 1398/4198 | 50 | 100 | 200 | 50 | 95 | 188 | 42 | 71 | 125 | 15 | 19 | 31 | 6 | 8 | 18 |
| 1RKE | 987/2947 | 40 | 83 | 163 | 50 | 100 | 199 | 25 | 41 | 73 | 18 | 30 | 49 | 39 | 61 | 107 |
| 1VG0 | 177/532 | 26 | 60 | 125 | 36 | 65 | 113 | 22 | 45 | 80 | 29 | 49 | 83 | 34 | 61 | 121 |
| 2B42 | 315/945 | 37 | 71 | 142 | 30 | 50 | 83 | 16 | 24 | 52 | 17 | 31 | 50 | 2 | 12 | 27 |
| 2HRK | 614/1842 | 8 | 29 | 69 | 43 | 83 | 160 | 35 | 53 | 93 | 25 | 43 | 79 | 49 | 95 | 177 |
| | Native/ Total | T10 | T20 | T40 | T10 | T20 | T40 | T10 | T20 | T40 | T10 | T20 | T40 | T10 | T20 | T40 |
| 2A9K | 31/94 | 3 | 9 | 18 | 7 | 12 | 22 | 6 | 11 | 17 | 5 | 10 | 16 | 8 | 13 | 25 |
| 2AYO | 40/120 | 6 | 6 | 11 | 10 | 18 | 31 | 1 | 4 | 12 | 1 | 2 | 8 | 4 | 10 | 22 |
| | Native/ Total | T5 | T10 | T20 | T5 | T10 | T20 | T5 | T10 | T20 | T5 | T10 | T20 | T5 | T10 | T20 |
| 1H1V | 9/29 | 2 | 4 | 7 | 2 | 4 | 7 | 2 | 3 | 7 | 1 | 2 | 6 | 0 | 0 | 3 |
| SR | - | 63% | 66% | 67% | 83% | 78% | 73% | 54% | 46% | 42% | 40% | 35% | 30% | 52% | 47% | 45% |

Next, we still counted the number of near native structures in the top N ranked docking decoys (N is 5, 10,20,40,50,100,200). In the Supplementary Table 2, Pisa achieved higher SR than SAGERank. This indicates that Pisa has a higher ranking accuracy for protein-protein docking decoys in the Top N. Nevertheless, SAGERank still has higher accuracy in ranking the top N of protein-protein docking decoys than the other three methods.

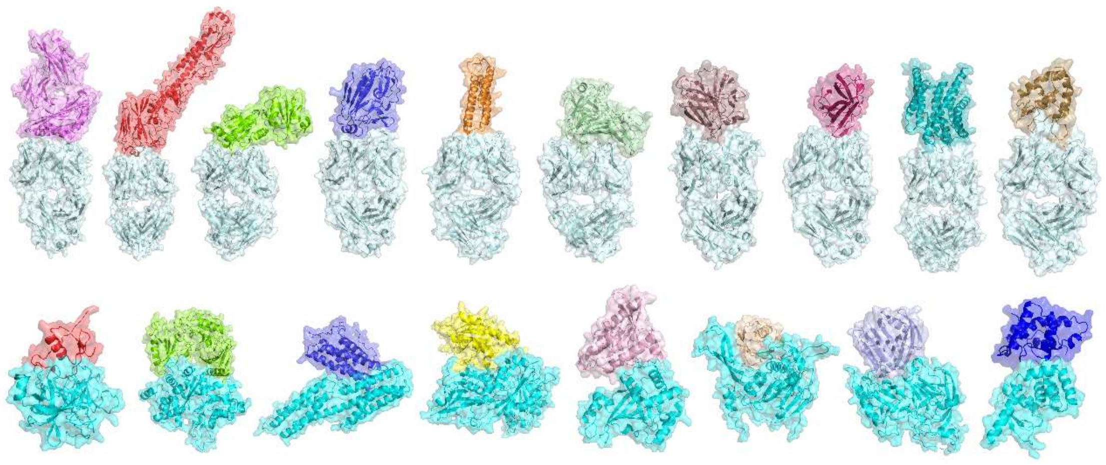

Supplementary Figure 5. The three-dimensional structure of antibody antigen complex (top) and protein protein complex (bottom) in the test set.

Supplementary Figure 5 is three-dimensional structure of 10 antibody-antigen complex and 8 protein-protein complex. Among them, dark blue and light blue structures serve as receptors in docking, while other colors serve as ligands. Similar to the processing method in the training set, semi flexible docking was used, with the receptor conformation unchanged, and the ligand conducts conformational search and energy optimization around the receptor's binding pocket or CDR region.

### Supplementary Note 2: Detecting antibody antigen binding

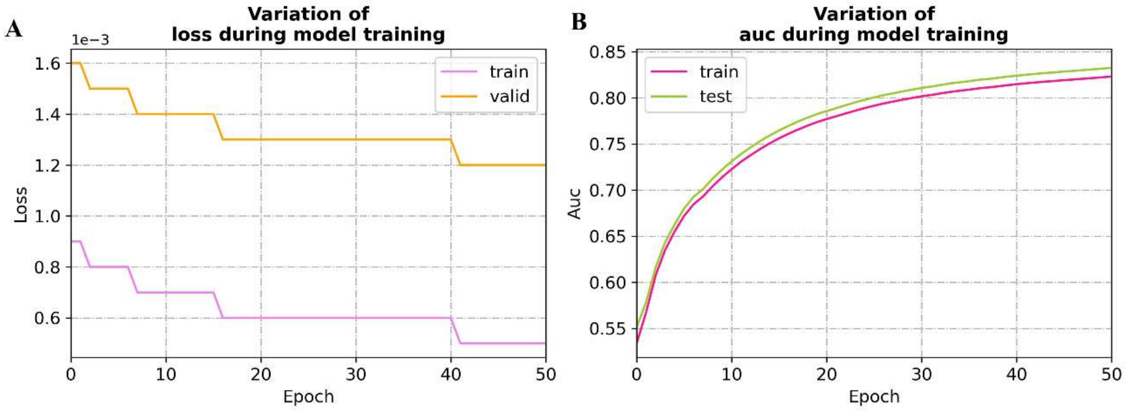


Supplementary Figure 6. The changes of loss (A) and AUC (B) during model training

Because the positive and negative samples in the test and training sets are not balanced (1:3), we used AUC instead of accuracy as an indicator here. From the picture 6B, it can be seen that the AUC of both the training and testing sets exceeds 0.8. This means that SAGERank can rank the majority of positive samples that antibody can bind to antigen before negative samples that antibody cannot bind to antigen.

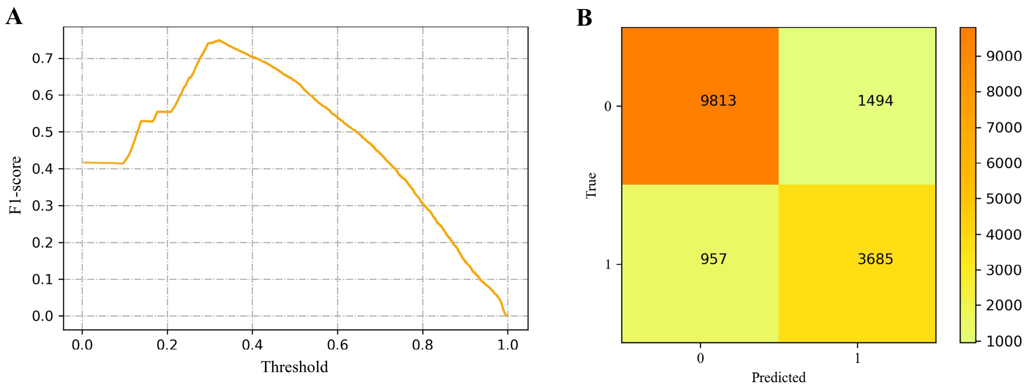


Supplementary Figure 7. F1-score curve with threshold variation (A) and confusion matrix of SAGERank on the test dataset (B)

In the F1-score curve with threshold variation (A), at a threshold of 0.3, F1 score can achieve a maximum value of 0.74. Therefore, choosing a threshold of 0.3 on the dataset for determining whether antibody antigen can be recognized can maximize the performance of the SAGERank. Figure 7B is confusion matrix of SAGERank on the test dataset that the threshold is 0.3. SAGERank accurately classified 9813 out of 11307 negative samples and 3685 out of 4642 positive samples.

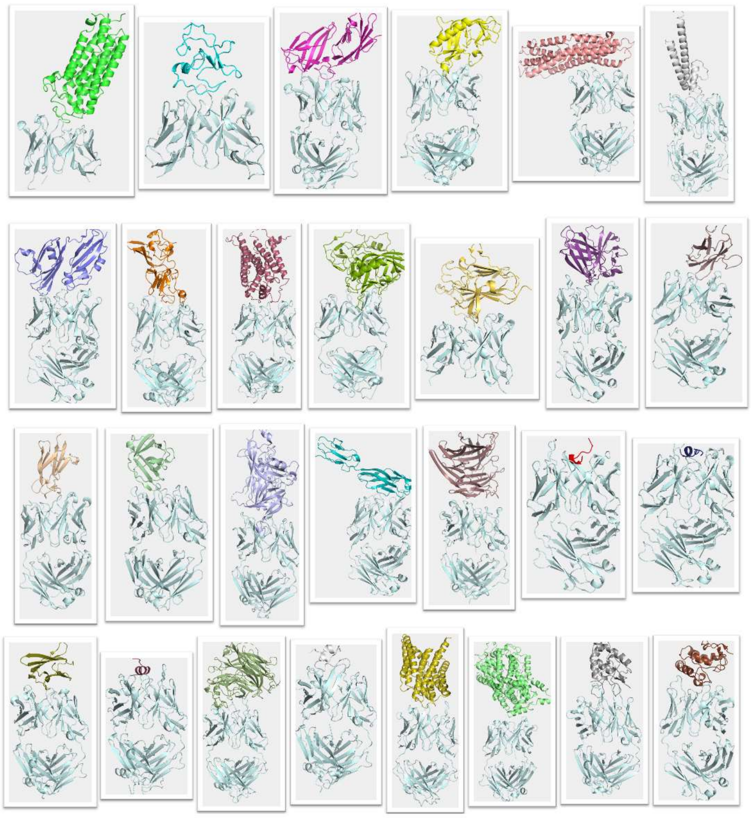

Supplementary Figure 8. The three-dimensional structure of antibody antigen complex in the test set

Figure 8 is a three-dimensional structural diagram of 30 different antibody antigen complexes in the test set. After docking with different antigens, a total of 15949 conformations were generated. Among them, there are 4642 antibodies that can bind to antigens (positive samples), and 11307 antibodies that cannot bind to antigens (negative samples).

### Supplementary Note 3: Identifying biological and crystallographic interfaces

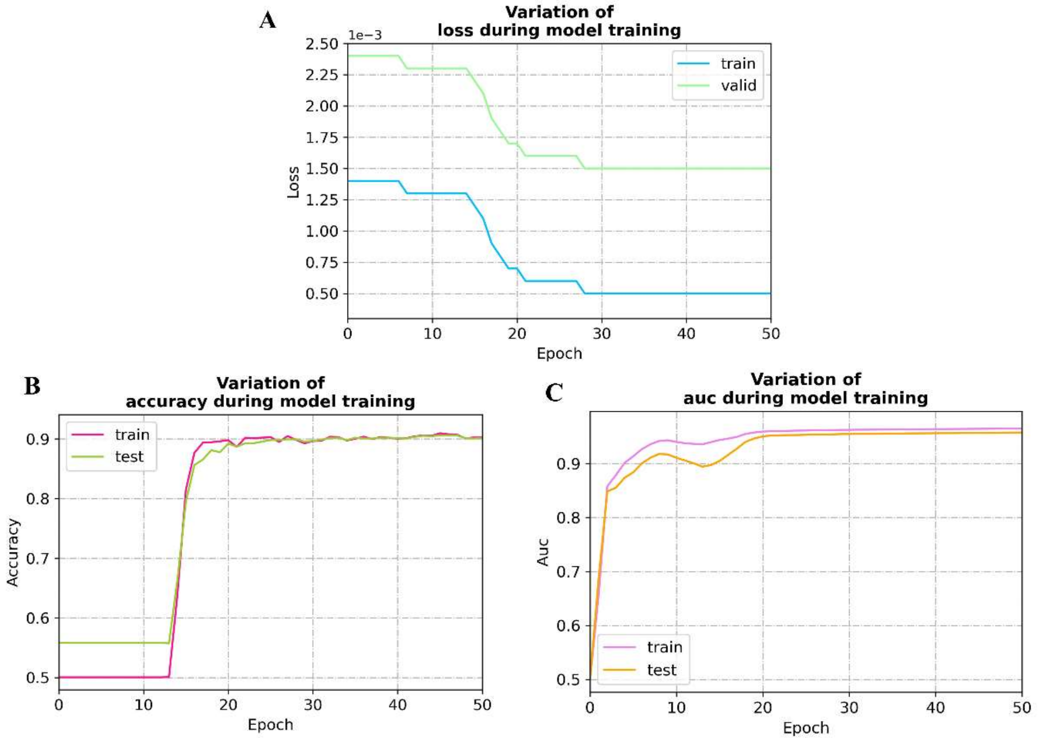


Supplementary Figure 9. The changes of loss (A), accuracy (B) and AUC (C) during model training

The Supplementary Figure 9 is the changes of loss, accuracy and AUC during model training, we can notice that the various indicators of the model perform very well and the accuracy and AUC reached 0.9 and 0.95, respectively.

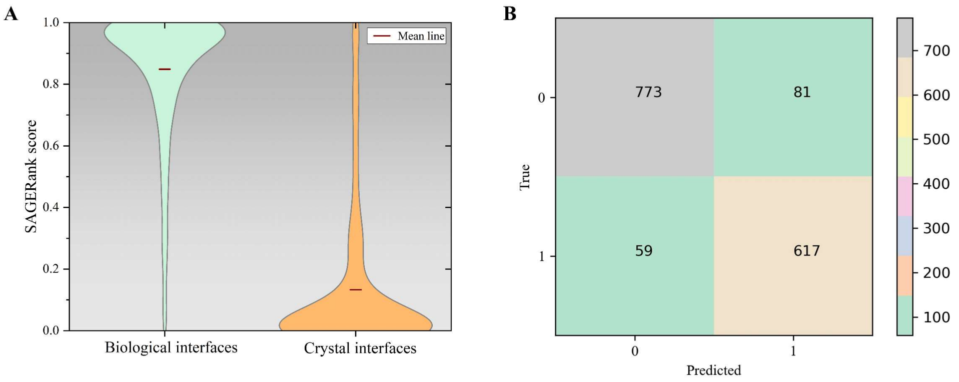


Supplementary Figure 10. The SAGERank scoring of biological and crystal interfaces in the test dataset (A) and confusion matrix of SAGERank on the test dataset (B)

From the Supplementary Figure 10A we can get that SAGERank shows a good polarization state in scoring the biological interface and crystal interface. The average scores for all biological interfaces and crystal interfaces are 0.845 and 0.13, respectively. In the Supplementary Figure 10B, SAGERank accurately classified 617 out of 698 biological interfaces (represented by 1) and 773 out of 832 crystal interfaces (represented by 0). This fully proves the classification accuracy of SAGERank.

### Supplementary Note 4: Some details during training for three applications

| Type | Application1 | Application2 | Application3 |
|---|---|---|---|
| Data size | 124G | 112G | 2.3G |
| Device | NVIDIA GeForce RTX 3090 | NVIDIA GeForce RTX 3090 | NVIDIA GeForce RTX 3080 |
| Training duration | 26days | 24days | 1.5days |
